# The role of gene expression on human sexual dimorphism: too early to call

**DOI:** 10.1101/2020.04.15.042986

**Authors:** Eleonora Porcu, Annique Claringbould, Kaido Lepik, BIOS Consortium, Tom G. Richardson, Federico A. Santoni, Lude Franke, Alexandre Reymond, Zoltán Kutalik

**Author notes:** These authors jointly supervised this work: Alexandre Reymond and Zoltán Kutalik. A full list of consortium members appears at the end of the paper (BIOS Consortium).

## Abstract

The genetic underpinning of sexual dimorphism is very poorly understood. The prevalence of many diseases differs between men and women, which could be in part caused by sex-specific genetic effects. Nevertheless, only a few published genome-wide association studies (GWAS) were performed separately in each sex. The reported enrichment of expression quantitative trait loci (eQTLs) among GWAS–associated SNPs suggests a potential role of sex-specific eQTLs in the sex-specific genetic mechanism underlying complex traits.

To explore this scenario, we performed a genome-wide analysis of sex-specific whole blood RNA-seq eQTLs from 3,447 individuals. Among 9 million SNP-gene pairs showing sex-combined associations, we found 18 genes with significant sex-specific *cis*-eQTLs (FDR 5%). Our phenome-wide association study of the 18 top sex-specific eQTLs on >700 traits unraveled that these eQTLs do not systematically translate into detectable sex-specific trait-associations. Power analyses using real eQTL- and causal effect sizes showed that millions of samples would be necessary to observe sex-specific trait associations that are fully driven by sex-specific *cis*-eQTLs. Compensatory effects may further hamper their detection. In line with this observation, we confirmed that the sex-specific trait-associations detected so far are not driven by sex-specific *cis*-eQTLs.

## Introduction

Men and women exhibit sexual dimorphism. Clear examples of sex-biased traits are anthropometric features. However, biological differences between sexes are not limited to physical traits: sex differences are also evident in incidence, prevalence and severity across diseases. For example, women are much more likely to develop autoimmune [1], while men are more likely to develop cardiovascular diseases [2].

Despite the widespread nature of these sexual differences and their noteworthy implications for medical research and treatments, little is known about their underlying biology in complex traits. While the sex chromosomes play key roles in sexual dimorphism, GWAS have identified dozens of autosomal genetic variants showing sex-specific effects [3–10], suggesting that part of the phenotypic differences might be due to accumulation of genetic variants present in both sexes at the same frequency [11], but acting in a different manner in males and females.

The strong enrichment of expression quantitative trait loci (eQTLs) among complex trait-associated loci [12–15] suggests that gene expression might be an appealing intermediate phenotype for the understanding of the biological mechanism behind SNP-trait associations. Towards this goal, several transcriptome-wide association studies (TWASs) integrating GWAS and eQTLs were proposed to identify genes whose expression is significantly associated to complex traits [16–18]. As these studies pointed to many genetic loci where variants exert their effect on phenotypes through gene expression, it is reasonable to think that sex-specific associations found by GWAS could be driven by sexual dimorphism in gene expression regulation, meaning that sex differences in eQTL effects might underlie the sex-specific GWAS associations. To explore this hypothesis, we performed a genome-wide investigation of sex biases in *cis*-eQTL effects in human autosomal genes and assessed their potential contribution to sex differences in complex traits.

## Results

### Sex-specific eQTLs analyses

First, we performed a genome-wide analysis of whole blood RNA-seq eQTLs to identify autosomal sex-specific eQTLs, i.e. SNPs whose effect on expression differs in magnitude between men and women. We analyzed eQTLs separately for 1,519 men and 1,928 women collected by the BIOS Consortium (http://www.bbmri.nl/acquisition-use-analyze/bios/). To reduce the number of tests, we restricted our analyses to autosomal variants previously detected as *cis*-eQTLs (FDR 5%) by the sex-combined analysis of the eQTLGen Consortium [19] and included in the UK10K reference panel [20].

To test the reliability of the BIOS data, we combined the results from the two sexes in a meta-analysis and compared them with those obtained by the eQTLGen Consortium (N=31,684, sex-combined results). We observed a high correlation between the betas obtained in the two studies (*r*^2^=0.9).

In total, we tested 8,739,806 SNP-gene pairs (involving 3,142,796 SNPs and 16,874 genes) for sex-interaction. We found 10,285 eGenes (genes with at least one *cis*-eQTL at Bonferroni adjusted *P*-value < 0.05) shared between the two sexes, while 542 and 1,419 eGenes were detected only in men or women, respectively.

### Identification of sex-specific cis-eQTLs

To identify sex-specific eQTLs, we tested the difference in the effects calculated for the two sexes separately (see Methods).

We identified 462 SNP-gene associations showing significantly different effects in men and women (FDR 5%, *P*_diff_<2.6×10^−06^). These sex-specific eQTLs cluster in 18 sex-specific eGenes (**Figure 1** and **Supplementary Table 1**). By analyzing only *cis*-eQTLs with a significant sex-combined effect (in the eQTLGen Consortium) we favored eQTLs that show the same direction but different magnitude of effect in men and women, as we posit that significant eQTLs showing opposite direction of effects in the two sexes would not have been detected in sex-combined analyses. Consistent with this hypothesis all 462 sex-specific eQTLs show the same direction but different magnitude of effects in both sexes.

**Figure1.**
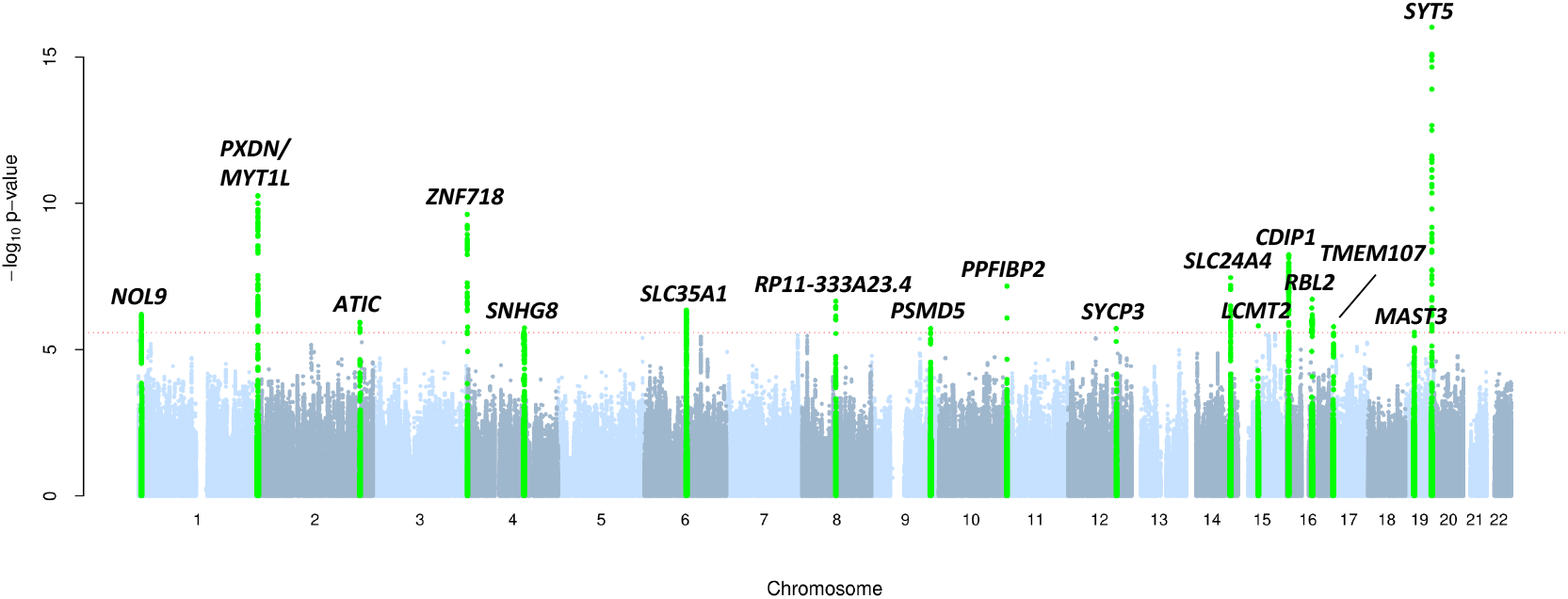
Manhattan plot of sex-specific eQTLs. The figure summarizes the results of our sex-specific eQTL discovery scan. SNP-gene pairs are plotted on the x-axis according to the SNP position on each autosomal chromosome in alternating light and dark blue against the *P*-values obtained upon testing for sex difference between effects in men and women (shown as −log_10_(*P*_diff_)). The red dotted line marks the 5% FDR threshold significance level (*P*_diff_=2.6×10^−6^), and SNPs in loci exceeding this threshold are highlighted in green.

Among the 18 sex-specific eGenes, 5 and 12 are men- and women-specific, respectively. One gene, *ZNF718*, contains eQTLs differently biased in both sexes (**Supplementary Figure 1**).

Next, to assess whether sex-specific eGenes are enriched for sex-specific biological processes, we performed enrichment analyses using 139 sex-specific eGenes, selected using a less stringent threshold (*P*_diff_ < 1×10^−4^). Although no GO term showed significant enrichment after FDR correction, we found response to glucagon (GO:0033762, *P*=1.59×10^−03^) and negative regulation of cytokine production (GO:1900016, *P*=2.43×10^−03^) as the two most enriched (top 50 GO terms are reported in **Supplementary Table 2**). These pathways are involved in two sexually dimorphic processes: glucose metabolism and inflammatory response, respectively [21] [22].

To determine if the difference of eQTL effects observed between men and women is driven by sex differences in gene expression distribution, we then compared the expression means and variances between sexes for the 18 sex-specific eGenes. While no eGene showed a significant difference in mean, we found 8 eGenes with sex-specific variance (based on *P*_diff_ < 0.05/18) (**Supplementary Table3**).

### Sex-specific cis-eQTLs do not translate into sex-specific trait associations

We then performed a phenome-wide association study (PheWAS) to test if eQTL SNPs with sex-specific effect on expression levels have an effect on human phenotypes and if so, whether sex biases in gene expression regulation translate to sex-specific effects on complex traits. For each sex-specific eGene we selected one representative eQTL with the strongest difference in effect between men and women and ran PheWAS analyses on more than 700 phenotypes from UK Biobank (UKBB) [23].

We found that 7 of the 18 lead eQTLs were associated with 39 traits at genome-wide significant level (**Supplementary Table 4**). Interestingly all associated traits belong to two categories: either morphological (e.g.: height, weight and trunk fat mass) or hematological traits (e.g.: platelets, eosinophils and whole blood cells).

We then asked if the other 290 non-lead sex-specific eQTLs of the 7 eGenes found by PheWAS showed also a sex-specific effect on the 39 pre-selected traits (we set a Bonferroni threshold of *P*_diff_<0.05/(20*7), where 20 is the effective number of independent phenotypes among the 39). For each trait, we used the sex-specific UKBB-GWAS summary statistics and observed no enrichment for the sex-specific eQTLs among the sex-specific GWAS signals and found only one eGene, *PSMD5*, for which sex-specific eQTLs are likely to translate to sex-specific associations with several obesity traits (**Figure 2**).

**Figure2.**
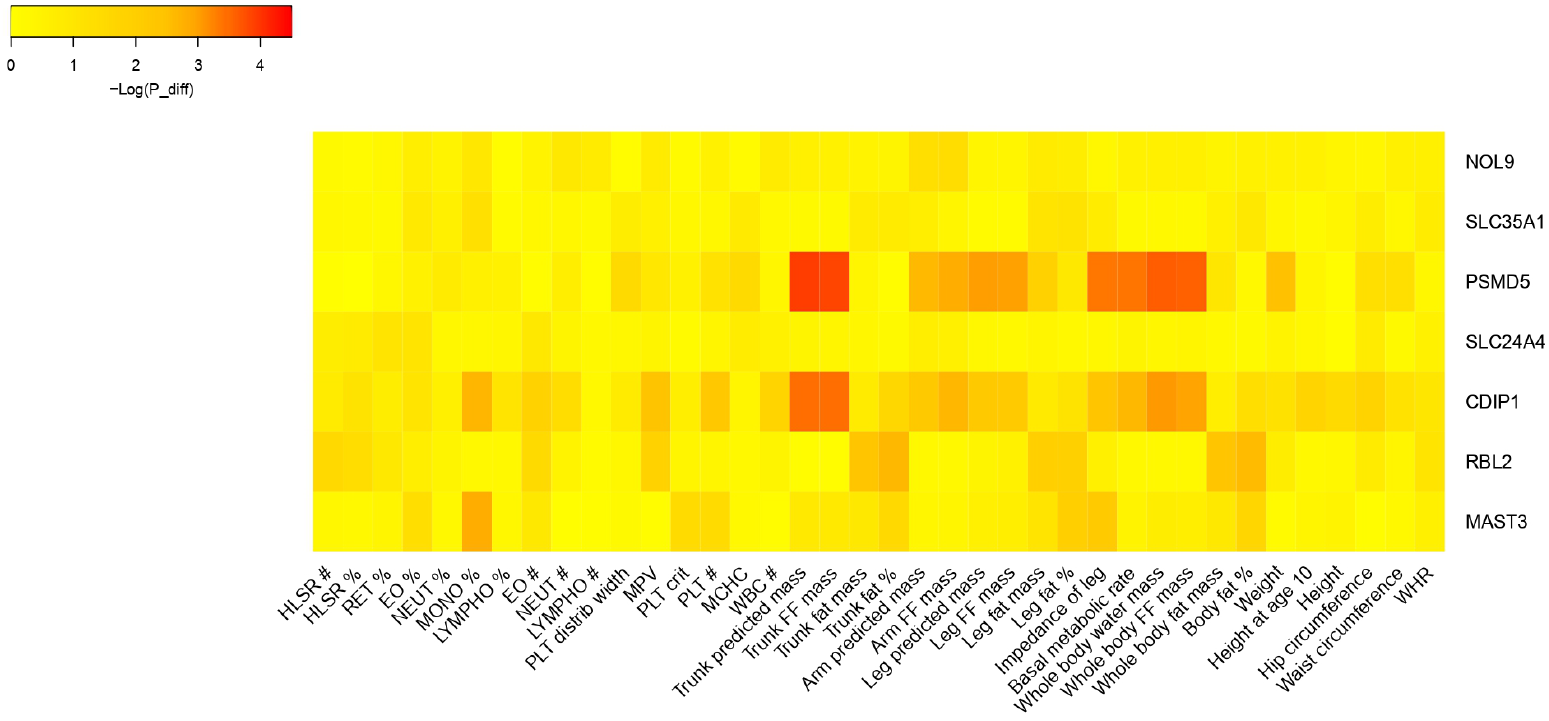
Heatmap of sex-specific trait-associated SNPs. The figure summarizes the results of sex-specific trait-associations driven by sex-specific eQTLs. For each sexspecific eGenes and for each phenotype, we plotted the minimum *P*_diff_ obtained testing for sex-difference between GWAS-effects in men and women (shown as −log_10_(*P*_diff_)).

### Sex-specific complex trait associations are not driven by sex-specific eQTLs

Next we tested whether sex-specific SNP-trait associations are driven by sex-specific gene expression regulation. For this we looked at the two most sexually dimorphic traits, waist-to-hip ratio (WHR) and testosterone levels. We identified 803 and 266 independent SNPs showing a *P*-value < 1×10^−05^ in the sex-combined GWAS for WHR and testosterone, respectively. Among those, 121 (32 for WHR and 89 for testosterone) have a significant sex-difference in the effect on the two sexes (*P*_diff_ <0.05/(803+266)), but none of the SNPs included in the BIOS Consortium dataset (58/121) show any sex-specific eQTL effect on genes in *cis* (*P*_diff_<0.05/58) (**Supplementary Tables 5 and 6**).

### Sex-specific causal effects

Next, to explore the presence of sex-specific causal effect of gene expression on complex traits, we performed sex-specific transcriptome-wide Mendelian Randomization (TWMR) analyses [18] combining sex-specific eQTLs and sexspecific GWAS results. We applied such approach – as above – to testosterone levels and WHR.

We found 17 and 6 genes showing a significantly different causal effect between the two sexes in testosterone and WHR, respectively (**Supplementary Figure 2 and 3** and **Supplementary Table 7**). While the sex-specific associations with WHR were all female-specific, among the 17 sex-specific genes associated with testosterone, 8 were female- and 9 male-specific, respectively.

Of note, the positive association of *IFT27* with testosterone levels observed only in men (*P*_TWMR-men_=2.55×10^−08^) and not in women (*P*_TWMR-women_=0.99) is supported by its association with Bardet-Biedl syndrome (OMIM: # 615996), for which it has been reported that most affected males produce reduced amounts of sex hormones (hypogonadism). Among the genes associated with WHR, *CCDC92* has a significant effect only in women (*P*_TWMR-women_=2.70×10^−23^) and not in men (*P*_TWMR-men_=0.09). Interestingly, diseases associated with *CCDC92* include lipodystrophy [24], which is known to affect more women than men [25].

As a negative control, we applied TWMR to a trait not showing sexual dimorphism, such as educational attainment, and did not observe any sex-specific gene association (**Supplementary Figure 4**).

Finally, we tested whether sex-specific causal effects are driven by sexspecific gene expression regulation and observed that none of the SNPs used as instrumental variables in TWMR showed a sex-specific effect on gene expression. In addition, we ran TWMR using sex-specific WHR GWAS and sex-combined eQTLs data and found a high correlation between the causal effects (*r^2^*=0.79 in females and *r^2^*=0.77 in males) (**Supplementary Figure 5**), which suggests that different effects observed by TWMR are driven by sex-specific SNP-trait associations.

### Power to detect sex-specific trait-associations

Since the observed sex-specific SNP-trait associations do not seem to be driven by sex-specific eQTL effects in our data, we performed power analyses using eQTL-effect differences observed in the data of the BIOS Consortium, which represents the largest biologically plausible sex-differences.

If the SNP-trait association is fully mediated by gene expression, then in females (F) and males (M)

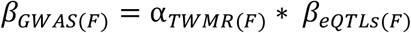

and

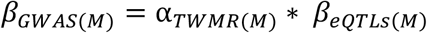

where *β_GWAS_* and *β_eQTLs_* indicate the effect of the SNP on the trait and on gene expression, respectively, and α_*TWMR*_ is the causal effect of the gene expression on the trait.

Assuming the same effect of the gene expression on the trait in both sexes (i.e.: α_*TWMR*(*M*)_ = α_*TWMR*(*F*)_ = α*_TWMR_*), the difference of SNP effect on the trait should be

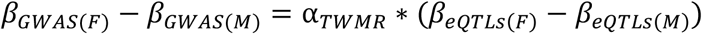

Using the differences observed by the BIOS Consortium and the causal effects estimated for a large number of complex traits by TWMR [18], we observed that the power to detect sex-specific trait-associations driven by sex-specific eQTLs is null, with an exception in the case when the largest causal effect of the gene expression on the trait observed in TWMR (100^th^ percentile) is coupled with the largest differences in eQTL-effects observed in BIOS Consortium (**Figure 3a**). We also estimated the GWAS sample size required to reach 80% power to detect differences in sex-specific GWAS. We found that, using the subset of unrelated British individuals from UKBB (N=380K) we do not have the power to detect sex-differences driven by SNPs being eQTLs for causal. Indeed, our results show that even when we used the 80^th^ percentile of the distribution of the significant causal effects of the gene expression on the trait, we need one to five million individuals to detect the different effect driven by sexspecific eQTLs (**Figure 3b**).

**Figure 3a.**
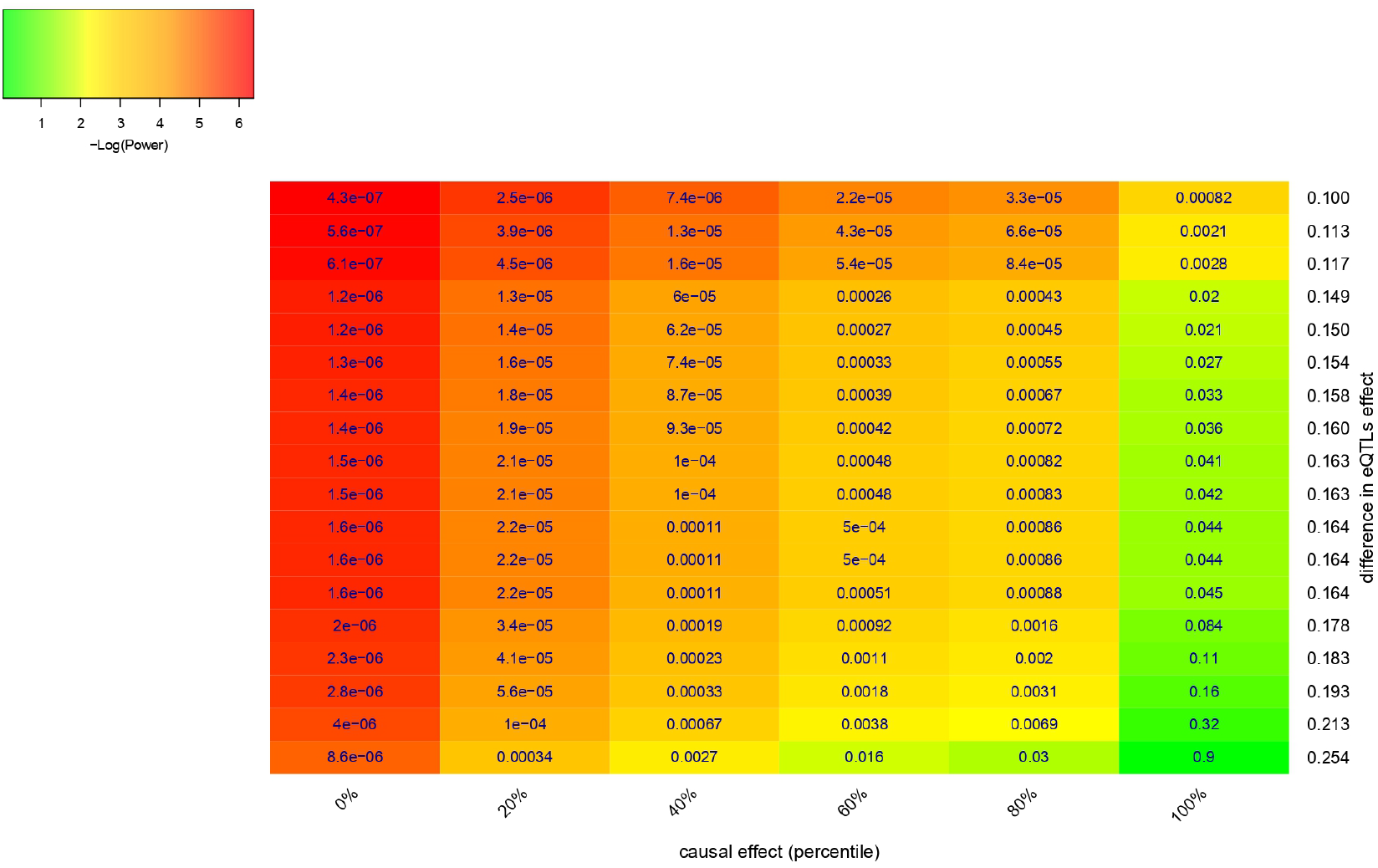
Heatmap showing the power to detect significantly different effect in sex-specific GWAS collecting 190,000 females and 170,000 males

**Figure 3b.**
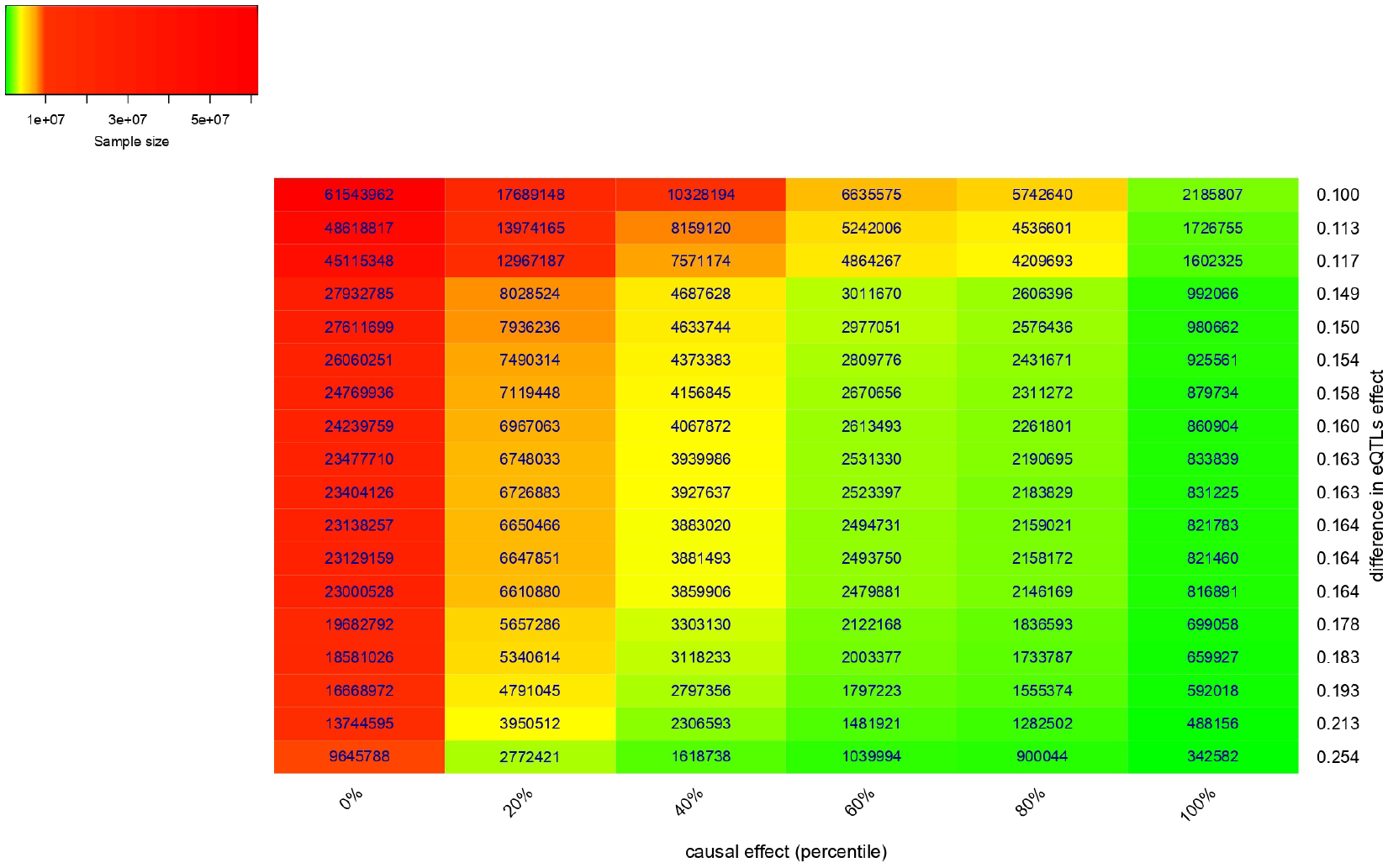
Heatmap showing the total sample size (females + males) needed to detect significantly different effect in sex-specific GWAS at power > 80%.

## Discussion

Notwithstanding the prominent differences in traits observed between men and women, there is little known about the role of sex-specific genetic effects. Several sex-stratified GWASs identified sex-specific genetic variants on autosomal chromosomes, which highlights the fact that not all differences are located on the sex chromosomes [3–10]. Since many genetic variants exert their effect on complex traits through gene expression [17, 18], sex differences in eQTL effects might underlie such sex-specific GWAS associations.

In this study, we used large sex-specific eQTL data and sex-specific GWAS results to investigate the role of gene expression on the sexual dimorphism of several human phenotypes.

We identified 18 sex-specific eGenes and 7 of them were associated with traits known to exhibit sex differences. Furthermore we found that the sex-specific eGenes are nominally enriched in GO biological processes linked to sex-specific processes.

Our results suggest that sex-specific eQTLs in whole blood do not translate to detectable sex-specific trait associations, and vice versa that the observed sex-specific trait associations cannot be explained by sex-specific eQTLs. While a recent work revealed that ~11% of trait heritability could be explained by *cis*-eQTL regulation [26], our findings show that the sex-specific heritability is not detectably mediated by sex-specific gene expression regulation. Our extensive power analyses, performed using a range of realistic effect sizes, confirmed these observations. Indeed we proved that with the current sample size used in sex-specific GWAS, we do not have the power to detect differences in sex-specific trait-associations driven by sex-specific eQTLs. Our results suggest that we will be able to explore how sex-specific gene expression regulation translates to complex traits only when GWAS will be performed on millions of individuals. It is only then that we will be able to test the existence of potential compensatory mechanisms via negative feedback loops dampening such signal propagation.

There are some limitations to this study. Firstly, although the BIOS RNA-seq dataset is relatively large (including 1,918 women and 1,519 men), it is limited to whole blood. Since it is known that the effect of causal genes on diseases typically act in a tissue-specific manner [27–29], the investigation of other, more relevant, tissues could be crucial to estimate larger causal effects and unravel the sex-specific associations found by the GWAS. Moreover, although we used the biggest sexspecific GWAS results, we convincingly show that the currently available sample size is too small to reach the statistical power necessary for detecting sex-specific trait-association mediated by sex-specific blood eQTLs.

As several studies have shown that eQTL effects can be cell type-specific [30, 31], upcoming single-cell eQTL datasets [32] might be essential in identifying sex- and cell-type specific effects and unravel the biological mechanism behind sexual dimorphism. Alternatively, if sexual dimorphism of complex traits is not driven by gene expression changes we might need to explore other types of omics data to gain a deeper insight into the molecular underpinnings of sex-differences in complex diseases.

## Methods

### Study sample

The Biobank-based Integrative Omics Study (BIOS, http://www.bbmri.nl/acquisition-use-analyze/bios/) Consortium has been set up in an effort of several Dutch biobanks to create a homogenized dataset with different levels of ‘omics’ data layers. Genotyping was performed in each cohort separately, as described before: LifeLines DEEP [33], Leiden Longevity Study [34, 35], Netherlands Twin Registry [36]; Rotterdam Study [37, 38], Prospective ALS Study Netherlands [39]. All genotypes were imputed to the Haplotype Reference Consortium [40] using the Michigan imputation server [41].

RNA-seq gene expression data was generated in The Human Genotyping facility (HugeF, Erasmus MC, Rotterdam, the Netherlands, http://www.blimdna.org). RNA-seq extraction and processing has been described before for a subset of the data [42]. Briefly, RNA was extracted from whole blood and paired-end sequenced using Illumina HiSeq 2000. Reads were aligned using STAR 2.3.0e [43] while masking common (MAF > 0.01) SNPs from the Genome of the Netherlands [44]. Gene-level expression was quantified using HTseq [45]. FastQC (http://www.bioinformatics.babraham.ac.uk/projects/fastqc/) was used to check quality metrics, and we removed individuals with < 70% of reads mapping to exons (exon mapped / genome). We included only unrelated individuals in this analysis and removed population outliers by filtering out samples with >3 standard deviations from the average heterogeneity score. We removed 25 PCs, from the expression matrix with all cohorts combined, to account for unmeasured variation.

We stratified the samples by sex and performed the *cis*-eQTL mapping using a pipeline described previously [46]. In brief, the pipeline takes a window of 1Mb upstream and 1Mb downstream around each SNP to select genes or expression probes to test, based on the center position of the gene or probe. The association between these SNP-gene combinations was calculated using a Spearman correlation in each sex separately.

### Differential gene expression and variance analysis

Differential expression analysis was preformed without considering genotypes, to identify genes with sex-specific expression (FDR 5%). For each gene, to test the difference in mean in the two sexes – calculated before any normalization but after quality check on samples – we used the t statistics

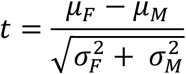

where *μ* and *σ* are the mean and the standard error, respectively.

To detect genes with sex-specific expression variance, we used normalized data and we applied two different strategies. First, we tested for difference in variance between females and males using an F-test, i.e.:

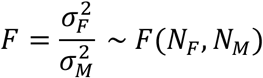

Second we used the t statistic

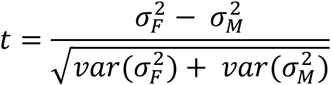

Genes showing significant *P*-values (FDR 5%) in both tests, were used for pathways analyses.

### Sex-specific eQTLs effects

To identify SNP-gene pairs with sex-difference, we computed *P*-values (*P*_diff_) testing for difference between the standardized men-specific and women-specific *β_eQTLs_*-estimates, with corresponding standard errors and using the t statistic

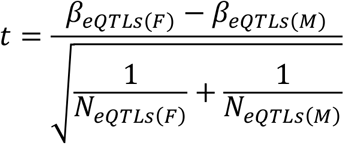

We selected 462 sex-specific SNP-gene pairs at an FDR of 5% across all the pairs tested.

### Sex-specific GWAS effects

To identify SNPs with sex-difference in the 39 associated phenotypes, we downloaded the summary statistics of the sex-stratified GWAS available at http://www.nealelab.is/uk-biobank/

We computed *P*-values (*P*_diff_) testing for a difference between the men-specific and women-specific *β_GWAs_*-estimates, with corresponding standard errors and using the t statistic

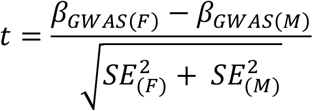

### Pathway analyses

In order to explore whether certain pathways are enriched among the genes containing eQTLs with evidence for sex-difference, we applied topGO. We tested the enrichment of GO terms with regard to the list of sex-specific eGenes at *P*_diff_<1×10^−4^ (list of genes including at least one eQTL showing *P*_diff_<1×10^−4^).

### Power analyses

We performed power analyses to calculate the probability that sex-specific SNPs found by GWAS are driven by sex-specific eQTLs. Using real observed data, we tested the power to detect a significant difference in *β*_GWAS_ in males and females starting from the difference observed in *β*_eQTLs_ and the causal effect of the gene expression on the phenotype calculated by TWMR (α_TWMR_).

If the association of a SNP in a given phenotype is driven by eQTLs, then we have

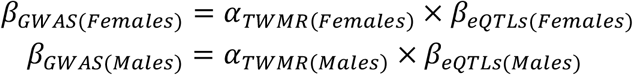

Let’s suppose that the effect of the gene expression on the phenotype is the same in the two sexes

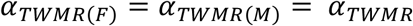

Then

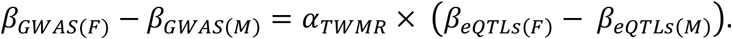

Since 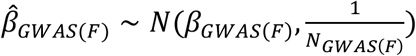 and 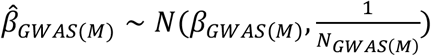, our statistics *t* follows a normal distribution when N is large,

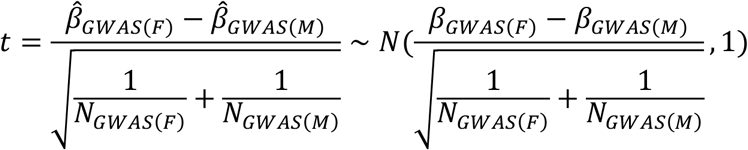

We tested the hypothesis 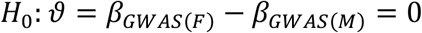 against 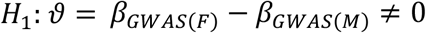.

Using the genome-wide significance threshold, *H*_0_ will be rejected if

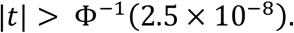

Then, the power to detect 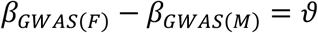 is

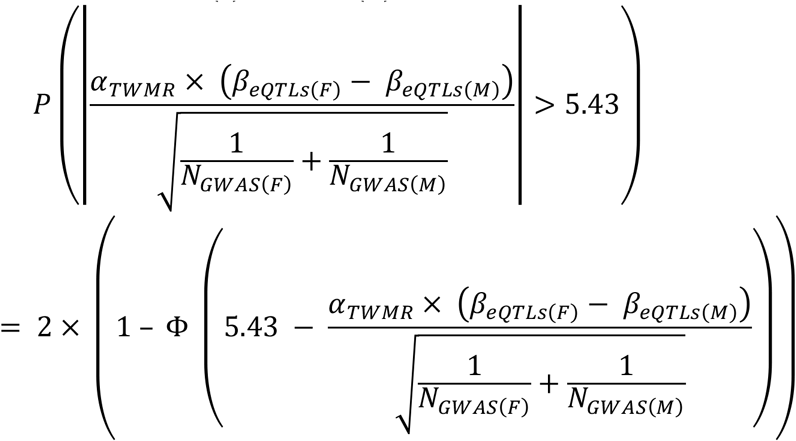

We performed the power analyses using the difference of *β_eQTLs_* observed in the 18 top sex-specific eQTLs and in 6 quantiles extracted from the distribution of significant causal effects estimated by TWMR for WHR.

Using the same statistics, we calculated the sample size to observe 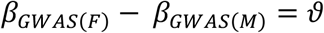 with power >0.8.

## Acknowledgments

This work was supported by grants from the Swiss National Science Foundation (31003A-143914 to ZK and 31003A_160203 to AR) and the Horizon2020 Twinning project ePerMed (692145 to AR). The funders had no role in study design, data collection and analysis, decision to publish, or preparation of the manuscript. We would like to thank Chiara Auwerx, Liza Darrous and Ninon Mounier for their valuable feedback and comments on this manuscript.

## Author Contributions

E.P., Z.K. and A.R. conceived and designed the study. E.P. performed statistical analyses. A.C. performed statistical analyses on BIOS Consortium dataset. L.F. supervised all statistical analyses performed in the BIOS Consortium. K.L. contributed by providing statistical support. E.P., T.G.R., F.A.S., L.F. A.R. and Z.K. contributed by providing advice on interpretation of results. E.P., A.R. and Z.K. wrote the manuscript with the participation of all authors. BIOS Consortium provided eQTLs data.

## BIOS Consortium (Biobank-based Integrative Omics Study) – Author information

**Management Team** Bastiaan T. Heijmans (chair)^1^, Peter A.C. ’t Hoen^2^, Joyce van Meurs^3^, Aaron Isaacs^4^, Rick Jansen^5^, Lude Franke^6^.

**Cohort collection** Dorret I. Boomsma^7^, René Pool^7^, Jenny van Dongen^7^, Jouke J. Hottenga^7^ (Netherlands Twin Register); Marleen MJ van Greevenbroek^8^, Coen D.A. Stehouwer^8^, Carla J.H. van der Kallen^8^, Casper G. Schalkwijk^8^ (Cohort study on Diabetes and Atherosclerosis Maastricht); Cisca Wijmenga^6^, Lude Franke^6^, Sasha Zhernakova^6^, Ettje F. Tigchelaar^6^ (LifeLines Deep); P. Eline Slagboom^1^, Marian Beekman^1^, Joris Deelen1, Diana van Heemst^9^ (Leiden Longevity Study); Jan H. Veldink^10^, Leonard H. van den Berg^10^ (Prospective ALS Study Netherlands); Cornelia M. van Duijn^4^, Bert A. Hofman^11^, Aaron Isaacs^4^, André G. Uitterlinden^3^ (Rotterdam Study).

**Data Generation** Joyce van Meurs (Chair)^3^, P. Mila Jhamai^3^, Michael Verbiest^3^, H. Eka D. Suchiman^1^, Marijn Verkerk^3^, Ruud van der Breggen^1^, Jeroen van Rooij^3^, Nico Lakenberg^1^.

**Data management and computational infrastructure** Hailiang Mei (Chair)^12^, Maarten van Iterson^1^, Michiel van Galen^2^, Jan Bot^13^, Dasha V. Zhernakova^6^, Rick Jansen^5^, Peter van ’t Hof1^2^, Patrick Deelen^6^, Irene Nooren^13^, Peter A.C. ’t Hoen^2^, Bastiaan T. Heijmans^1^, Matthijs Moed^1^.

**Data Analysis Group** Lude Franke (Co-Chair)^6^, Martijn Vermaat^2^, Dasha V. Zhernakova^6^, René Luijk^1^, Marc Jan Bonder^6^, Maarten van Iterson^1^, Patrick Deelen^6^, Freerk van Dijk^14^, Michiel van Galen^2^, Wibowo Arindrarto^12^, Szymon M. Kielbasa^15^, Morris A. Swertz^14^, Erik. W van Zwet^15^, Rick Jansen^5^, Peter-Bram ’t Hoen (Co-Chair)^2^, Bastiaan T. Heijmans (Co-Chair)^1^.

^1^ Molecular Epidemiology Section, Department of Medical Statistics and Bioinformatics, Leiden University Medical Center, Leiden, The Netherlands

^2^ Department of Human Genetics, Leiden University Medical Center, Leiden, The Netherlands

^3^ Department of Internal Medicine, ErasmusMC, Rotterdam, The Netherlands

^4^ Department of Genetic Epidemiology, ErasmusMC, Rotterdam, The Netherlands

^5^ Department of Psychiatry, VU University Medical Center, Neuroscience Campus Amsterdam, Amsterdam, The Netherlands

^6^ Department of Genetics, University of Groningen, University Medical Centre Groningen, Groningen, The Netherlands

^7^ Department of Biological Psychology, VU University Amsterdam, Neuroscience Campus Amsterdam, Amsterdam, The Netherlands

^8^ Department of Internal Medicine and School for Cardiovascular Diseases (CARIM), Maastricht University Medical Center, Maastricht, The Netherlands

^9^ Department of Gerontology and Geriatrics, Leiden University Medical Center, Leiden, The Netherlands

^10^ Department of Neurology, Brain Center Rudolf Magnus, University Medical Center Utrecht, Utrecht, The Netherlands

^11^ Department of Epidemiology, ErasmusMC, Rotterdam, The Netherlands

^12^ Sequence Analysis Support Core, Leiden University Medical Center, Leiden, The Netherlands 13. SURFsara, Amsterdam, the Netherlands

^14^ Genomics Coordination Center, University Medical Center Groningen, University of Groningen, Groningen, the Netherlands

^15^ Medical Statistics Section, Department of Medical Statistics and Bioinformatics, Leiden University Medical Center, Leiden, The Netherlands

