## Supplementary Figures 2_5 for "The role of gene expression on human sexual dimorphism: too early to call"

**Supplementary Figure 2.** Miami plot for testosterone. SNPs are plotted on the x-axis according to their position on each autosomal chromosome against the P-values (shown as –/+ $\log_{10}$(P-value)) obtained upon testing for association in women (pink dots) and men (blue dots). Loci containing sex-specific causal genes are highlighted in green.

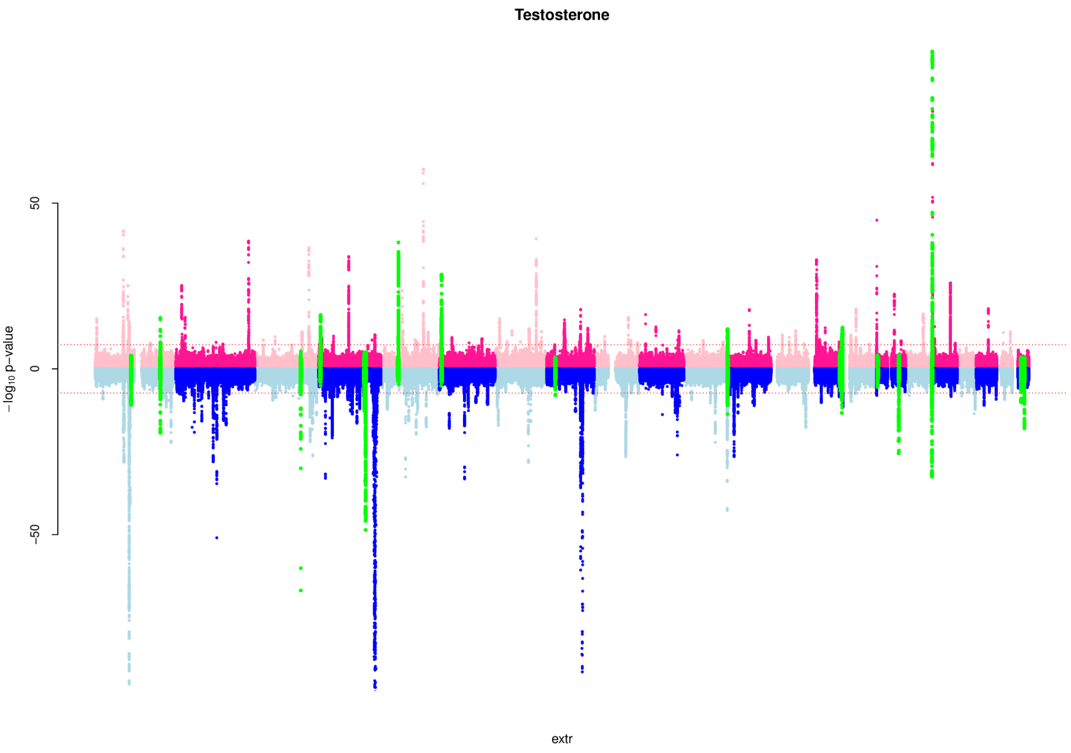

**Supplementary Figure 3.** Miami plot for waist-to-hip ratio (WHR). SNPs are plotted on the x-axis according to their position on each autosomal chromosome against the P-values (shown as –/+ $log_{10}$(P-value)) obtained upon testing for association in women (pink dots) and men (blue dots). Loci containing sex-specific causal genes are highlighted in green.

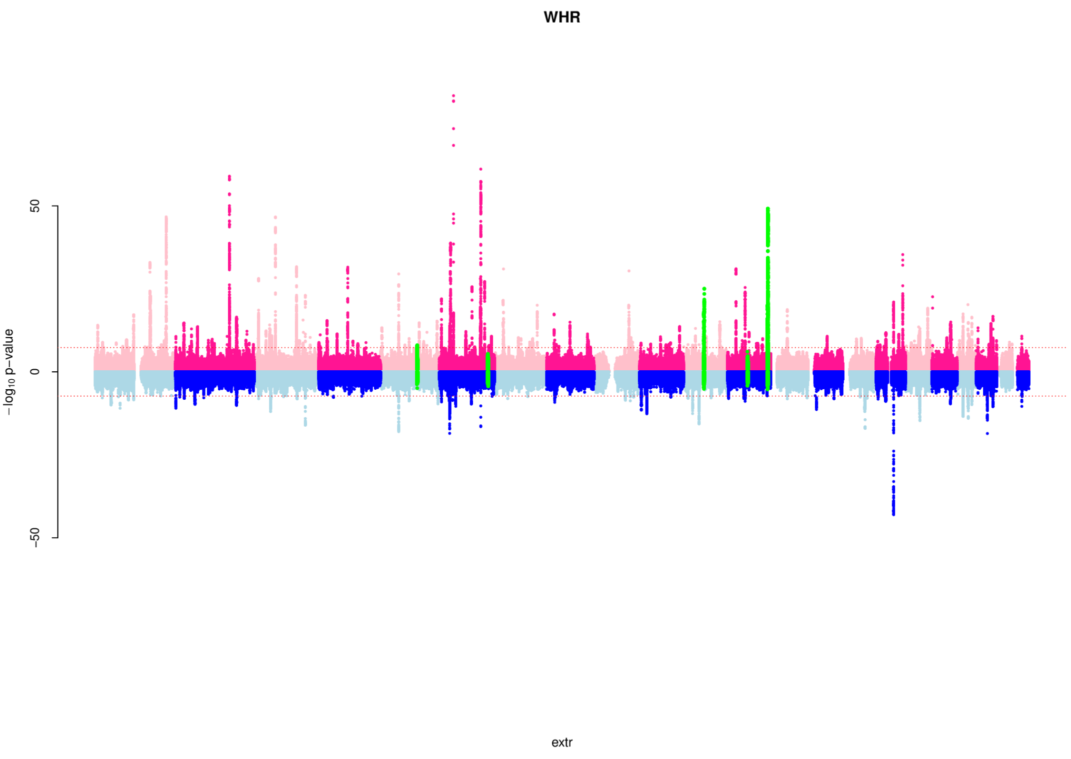

**Supplementary Figure 4.** Miami plot for educational attainment. SNPs are plotted on the x-axis according to their position on each autosomal chromosome against the P-values (shown as –/+ $\log_{10}$(P-value)) obtained upon testing for association in women (pink dots) and men (blue dots).

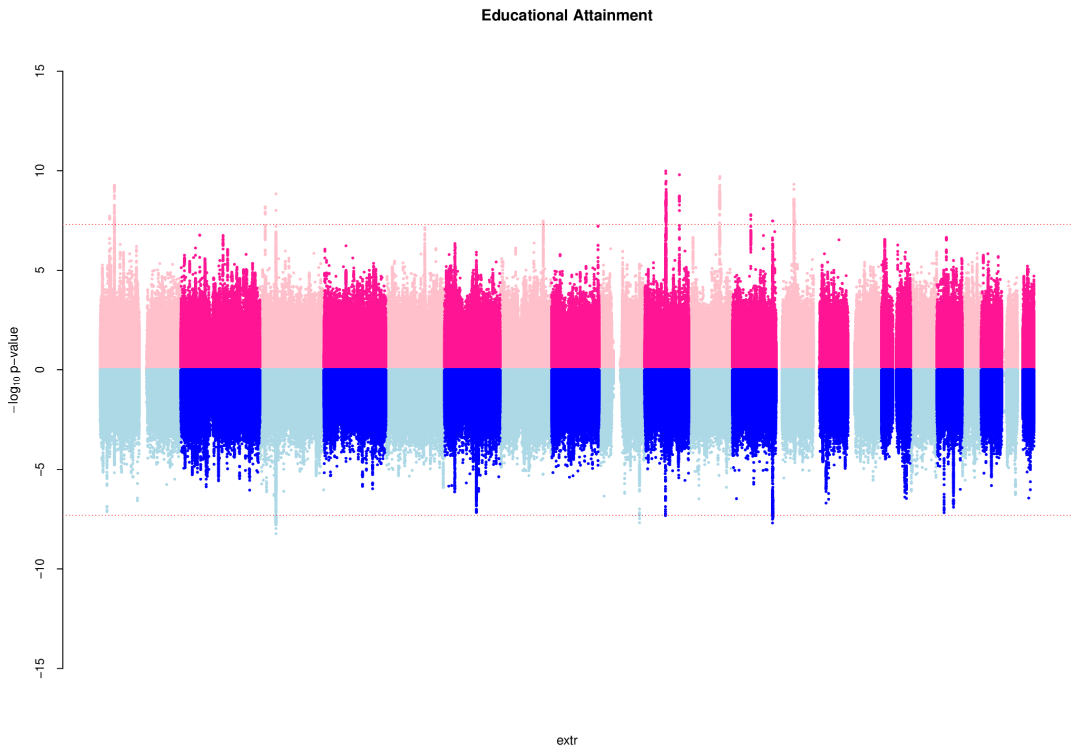


**Supplementary Figure 5.** Comparison of TWMR-causal effects estimated for WHR using sex-specific and combined eQTLs data.

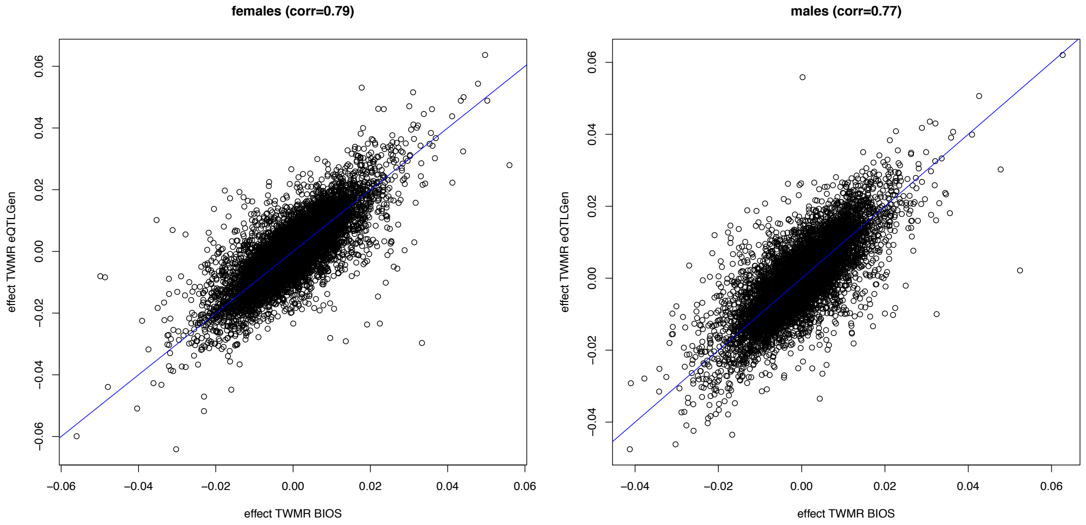
