## Supplementary Figure 1 for "The role of gene expression on human sexual dimorphism: too early to call"

**Supplementary Figure 1**. Comparison between the effects in the two sexes of the eQTLs in the 18 sex-specific eGenes. eQTLs showing a significant different effect in males and females are highlighted in red.

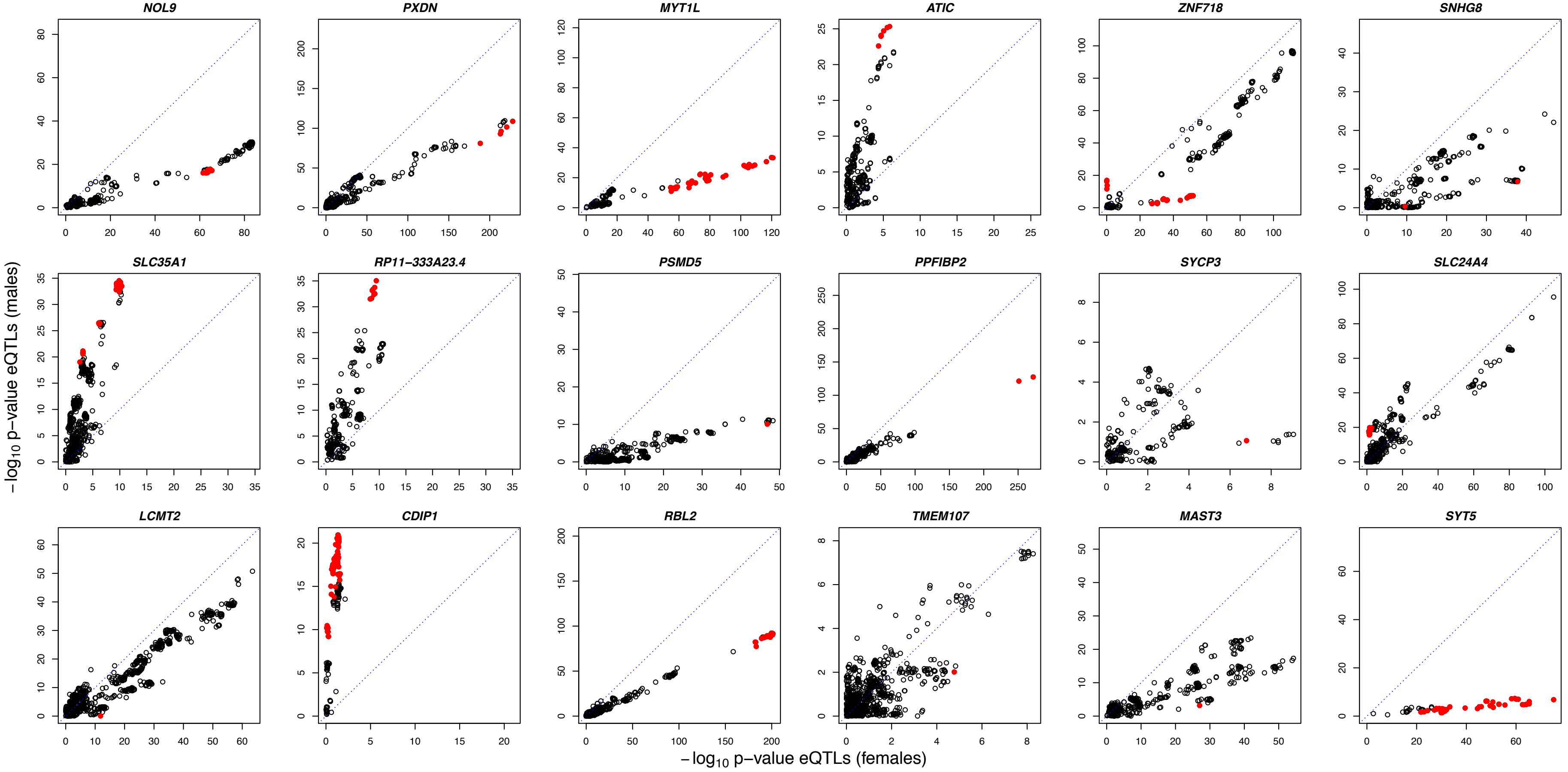
